# Oncogenic signals prime cancer cells for toxic cell growth during a G1 cell cycle arrest

**DOI:** 10.1101/2022.09.08.506962

**Authors:** Reece Foy, Lisa Crozier, Aanchal U Pareri, Ben Ho Park, Adrian T Saurin

## Abstract

A long-term goal in cancer research has been to inhibit the cell cycle in tumour cells without causing toxicity in proliferative healthy tissues. The best evidence that this is achievable is provided by CDK4/6 inhibitors, which arrest the cell cycle in G1, are well-tolerated in patients, and are effective in treating ER+/HER2-breast cancer. CDK4/6 inhibitors are effective because they arrest tumour cells more efficiently than some healthy cell types and, in addition, they affect the tumour microenvironment to enhance anti-tumour immunity. We demonstrate here another reason to explain their efficacy. Tumour cells are specifically vulnerable to CDK4/6 inhibition because during the G1 arrest, oncogenic signals drive toxic cell overgrowth. This overgrowth causes permanent cell cycle withdrawal by either preventing exit from G1 or by inducing replication stress and genotoxic damage during the subsequent S-phase and mitosis. Inhibiting or reverting oncogenic signals that converge onto mTOR can rescue this excessive growth, DNA damage and cell cycle exit in cancer cells. Conversely, inducing oncogenic signals in non-transformed cells can drive these toxic phenotypes and sensitize cells to CDK4/6 inhibition. Together, this demonstrates how oncogenic signals that have evolved to stimulate constitutive tumour growth and proliferation can be driven to cause toxic cell growth and irreversible cell cycle exit when proliferation is halted in G1.

## INTRODUCTION

Identifying cell cycle vulnerabilities that distinguish cancer cells from healthy cells has been a long-term goal in cancer research (Liu et al., 2022; Suski et al., 2021). A major breakthrough in this area came with the development of CDK4/6 inhibitors, which have revolutionised the treatment of the main subtype of metastatic breast cancer (ER+/HER2-) by increasing progression-free and overall survival when used in combination with anti-estrogen therapy (Fassl et al., 2022). The rationale for this combination is that blocking estrogen signalling inhibits the transcription of Cyclin D, the regulatory subunit of CDK4/6, thus producing a “double-hit” on Cyclin D-CDK4/6 activity, specifically in breast cancer cells that overexpress estrogen receptors. This leads to an efficient G1 arrest, because Cyclin D-CDK4/6 is required to phosphorylate Rb and thereby activate E2F transcription factors, which induce the expression of many genes required for S-phase (Kent and Leone, 2019). Oncogenic signals also act to drive excessive Cyclin D production in many tumour types, including breast cancer, and this has rationalised ongoing clinical trials to test whether inhibiting these signals alongside CDK4/6 can produce a similar double-hit to efficiently and specifically arrest tumour cell proliferation (Alvarez-Fernandez and Malumbres, 2020; Fassl et al., 2022). In addition to being particularly sensitive to these drug combinations, tumours are also thought to be intrinsically sensitive to CDK4/6 inhibition because tumour cells rely on the Cyclin D-CDK4/6 pathway for G1 progression more than some healthy cell types (Choi et al., 2012; Gong et al., 2017). This is likely due to constant stimulation of the pathway by either oncogenic signals, overexpression of Cyclin D/CDK4/CDK6, or loss/inhibition of tumour suppressors that restrain CDK4/6 activity (p16^INK4A^ and p53/p21) (Kent and Leone, 2019).

A crucial issue in the context of anti-cancer therapy concerns not just how efficiently tumour cells arrest in G1, but how cells then respond to that arrest. A positive response is often associated with marked tumour regression that is sustained well after therapy has ceased, implying that tumour cells experience a cytotoxic response to these drugs and not simply a cytostatic arrest. There are many ideas surrounding why this could occur, including that CDK4/6 inhibitors have intrinsic effects on tumour metabolism and extrinsic effects on the surrounding microenvironment to enhance anti-tumour immunity (Klein et al., 2018). However, a major gap in our understanding concerns the questions of when, why and how a pause in G1 transitions into a state of irreversible cell cycle exit, known as senescence (Wagner and Gil, 2020). It is critical to address these questions because it may help to explain why tumours are more sensitive to these drugs than healthy cells, and this may ultimately help us to better predict the most sensitive tumour types and/or the best drug combinations. We therefore set out to resolve these issues by building on our recent data demonstrating that a pause in G1, if held for too long, downregulates various replisome components to cause DNA damage and long-term cell cycle withdrawal after release from that arrest (Crozier et al., 2022b). The crucial question we sought to address was: what happens during the G1 arrest that causes such widescale proteomic changes that ultimately leads to problems during the subsequent cell cycle?

## RESULTS

### Cellular overgrowth following CDK4/6 inhibition causes genotoxic stress and cell cycle exit

One clear effect of pausing cells for long periods in G1 is that they become progressively enlarged in size, as total cellular protein and RNA continue to increase despite the cell cycle arrest. This occurs in non-transformed hTERT-RPE1 cells (RPE1) and in ER+/HER2-breast cancer cells that are p53-proficient (MCF7) or p53-deficient (T47D) (**Figure 1A-C;** and see accompanying paper by Crozier et al). This is broadly consistent with similar observations reported recently by others (Ginzberg et al., 2018; Lengefeld et al., 2021; Neurohr et al., 2019; Zatulovskiy et al., 2020). We sought to prevent this excessive growth during G1, so that we could test whether it was responsible for the downstream effects on DNA damage and long-term cell cycle exit. Excessive growth during G1 was PI3K/mTOR-dependent because it was completely prevented by co-treatment with PF-05212384; a dual inhibitor of PI3K and mTOR (Mallon et al., 2011) (**Figure 1C-D and S1A-B**). FUCCI analysis demonstrates that when RPE1 cells are released from a prolonged G1 arrest they struggle to renter the cell cycle, and those cells that do enter S-phase frequently revert back into G1 without entering mitosis (**Figure 1E**), as demonstrated recently (Crozier et al., 2022b). Importantly, both of these phenotypes were rescued when overgrowth was prevented with PF-05212384 (**Figure 1E-F**), which was associated with a dramatic improvement in long-term proliferation (**Figure 1G**). Cell cycle re-entry and progression were similarly improved in breast cancer cells following PF-05212384 treatment (**Figure S1C-D**).

**Figure 1:**
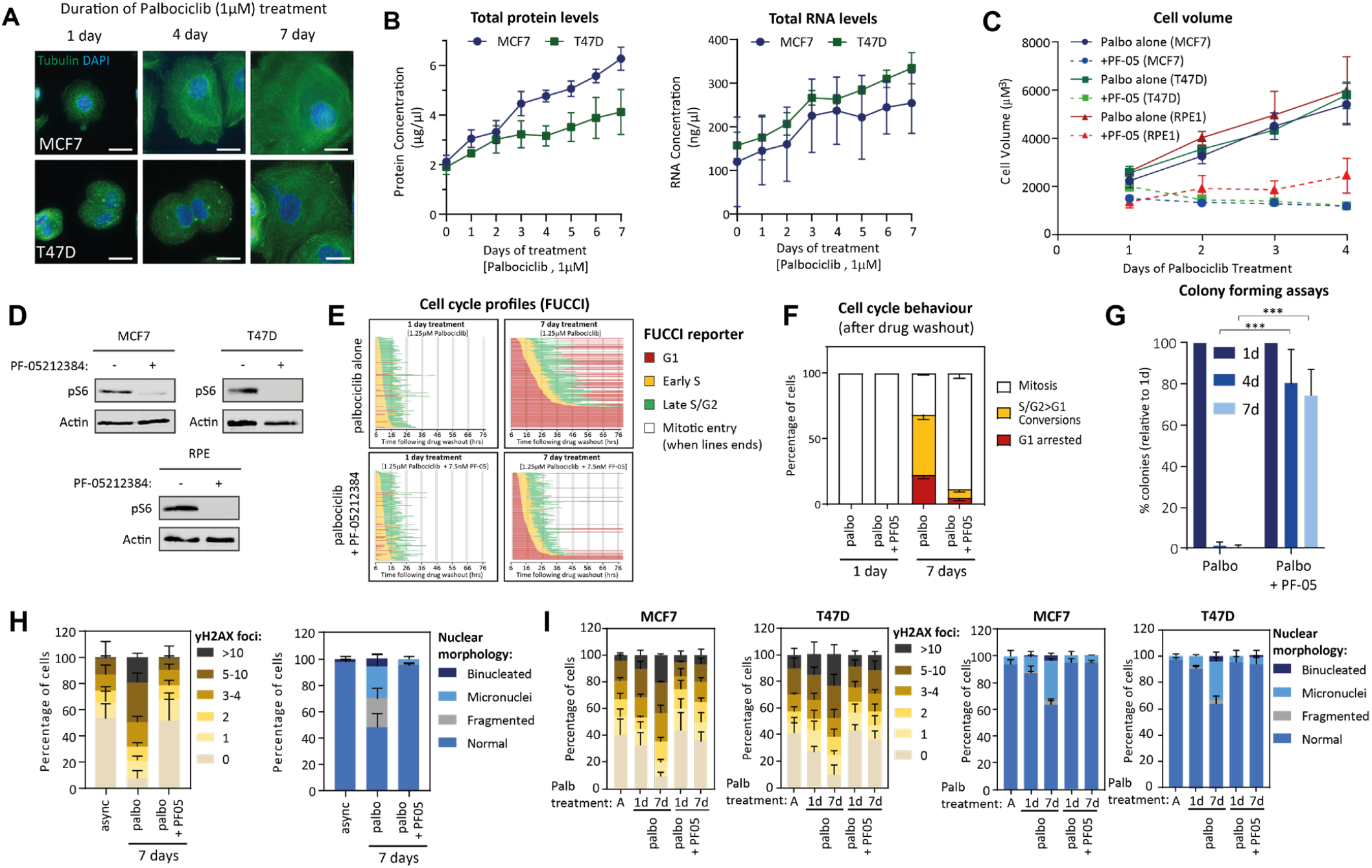
mTOR-dependent overgrowth during a G1 arrest drives DNA damage and cell cycle exit. **A-B)** Immunofluorescence images (A), and protein and RNA concentration measurements (B) of MCF7 and T47D cells arrested in palbociclib (1μM) for 0-7 days. The channel intensities in A are scaled differently between conditions to prevent tubulin saturation in small cells. Scale bars = 25μM. **C)** Cell volume assays of cells arrested in palbociclib (1.25uM for RPE, 1μM for MCF7/T47D) for 1-4 days in the presence/absence of PF-05212384 (30nM for RPE1, 7.5nM for MCF7/T47D; see Figures S1A-B for dose response). Graph shows mean data -/+ SD from three repeats. **D)** Western analysis of cells arrested in palbociclib (1.25uM for RPE, 1μM for MCF7/T47D) for 1 days in the presence/absence of PF-05212384 (30nM for RPE1, 7.5nM for MCF7/T47D). Representative example of at least 4 repeats. **E)** Cell cycle profile of individual RPE1-FUCCI cells (each bar represents one cell) after washout from 1 or 7 days of treatment with palbociclib (1.25 μM) (Figure 1: continued) +/- PF-05212384 (30nM). STLC (10 μM) was added to prevent progression past the first mitosis. Fifty cells were analysed at random for each repeat and three experimental repeats are displayed (150 cells total). **F)** Quantifications of cell cycle defects from the single-cell profile plots displayed in E. Bar graphs show mean - SD. **G)** Colony forming assays in RPE1 cells treated with palbociclib (1.25 μM) +/- PF-05212384 (30nM) for 1, 4 or 7 days and then grown at low density without inhibitor for 10 days. Each bar displays mean data + SD from three experiments. Statistical significance determined by Fisher’s exact test (*** p < 0.0001). **H-I)** Quantification of γH2AX-positive foci (left panel) and nuclear morphologies (right panel) following palbociclib treatment in p53-KO RPE1 cells (H) or MCF7/T47D cells (I). Cells were treated with DMSO (asynch) or palbociclib for 1 or 7 days in the absence/presence of PF-05212384, as indicated, and then analysed after drug washout for 48 h (RPE1 cells) or 72h (MCF7/T47D). Palbociclib was used at 1.25 μM in RPE1 cells and 1 μM in MCF7/T47D, PF-05212384 was used at 30nM in RPE1 and 7.5nM in MCF7/T47D. A total of 100 cells (nuclear morphology) or 50 cells (γH2AX foci) were scored per condition per experiment, and bar graphs represent mean data + SD from three experiments.

Irreversible cell cycle withdrawal following CDK4/6 inhibition has recently been linked to replication stress as a result of impaired origin licencing and the progressive downregulation of replisome components during the G1 arrest (Crozier et al., 2022b). This replication stress induces p53-dependent cell cycle exit from G2, or in the absence of p53, excessive DNA damage during mitosis as the under-replicated chromosomes are mis-segregated to produce γH2AX foci and gross nuclear abnormalities. In agreement with a crucial rule for cell overgrowth in these phenotypes, PF-05212384 was able to rescue γH2AX foci and nuclear abnormalities following release from prolonged CDK4/6 inhibition in p53-KO RPE1 cells (**Figure 1H**), which typically have the highest rates of damage (Crozier et al., 2022b)), and in breast cancer cells that are p53-proficient (MCF7) or deficient (T47D) (**Figure 1I**).

In summary, excessive cell growth during a G1 arrest drives permanent cell cycle exit by restricting exit from G1 and by causing DNA damage in those cells that do re-enter the cell cycle. Inhibiting PI3K/mTOR signalling can completely prevent this growth, DNA damage and long-term cell cycle exit, thus explaining why mTOR activity is crucial to drive quiescent G1-arrested cells into senescence (Leontieva and Blagosklonny, 2013; Leontieva et al., 2013; Maskey et al., 2021). Therefore, mTOR status critically determines whether the arrest following CDK4/6 inhibition is cytotoxic or cytostatic.

In an accompanying paper, Crozier et al. explain mechanistically how proteome remodelling and osmotic stress in overgrown cells leads to downstream effects that culminate in prolonged cell cycle exit (Crozier et al., 2022a). Here we seek to examine the upstream signals that drive this overgrowth, and to explore whether these vary between cell types, since this would be predicted to have a crucial effect on outcome following CDK4/6 inhibition.

### Cancer cells are sensitised to overgrowth and cell cycle exit following CDK4/6 inhibition

The level of cell growth and mTOR activity is determined by the balance of growth promoting and growth repressing signals, which importantly, depends on both cell context and cell type. In non-transformed epithelial cells, growth factor signalling stimulates mTOR to drive cell growth and proliferation, however, upon cell-cell contact these signals are rapidly shut-down by contact inhibition of proliferation (Mendonsa et al., 2018). **Figure 2A-E** demonstrates that both serum withdrawal and cell-cell contact can inhibit mTOR and protect RPE1 cells from toxic overgrowth during a G1 arrest, thereby limiting cell cycle exit and restoring long-term proliferation following release from that arrest. This is in sharp contrast to breast cancer cells which continue to activate mTOR and grow during a G1 arrest, despite culturing in low serum or at high confluence (**Figure 2F-G**). Therefore, two pervasive hallmarks of cancer – loss of contact inhibition and growth-factor independence – facilitate the overgrowth of cancer cells following CDK4/6 inhibition.

**Figure 2:**
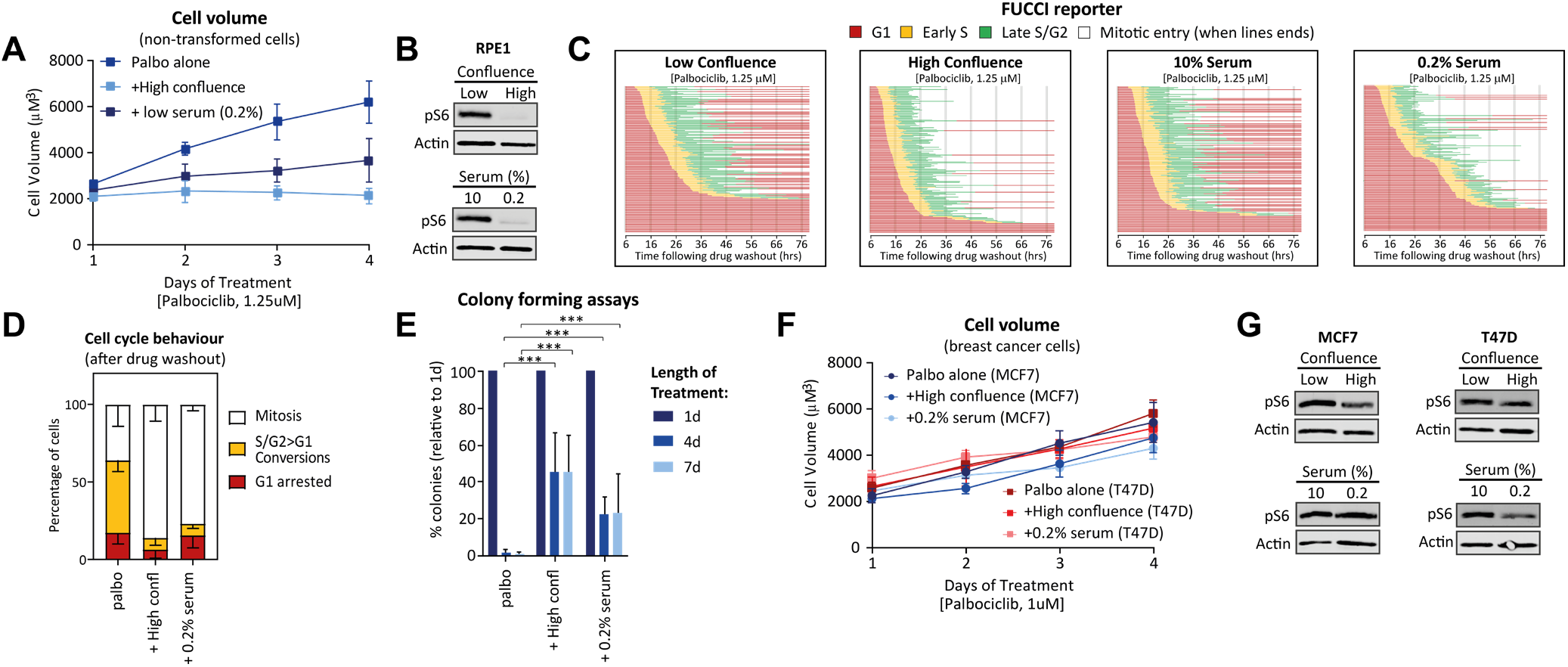
Serum withdrawal and contact inhibition can protect non-transformed cells from overgrowth and cell cycle exit following CDK4/6 inhibition. **A)** Cell volume assays of RPE1 cells arrested in palbociclib (1.25μM) for 1-4 days under control conditions (10% serum and low confluence), in low (0.2%) serum, or at high confluence. Graph shows mean data -/+ SD from three repeats. **B)** Western analysis of RPE1 cells arrested in palbociclib (1.25μM) for 1 day, but treated as in A. Representative example of at least 4 repeats. **C)** Cell cycle profile of individual RPE1-FUCCI cells (each bar represents one cell) after washout from 7 days of treatment with palbociclib (1.25 μM). During treatment cells were plated at high confluence or with low serum (0.2% FBS). STLC (10 μM) was added to prevent progression past the first mitosis. Fifty cells were analysed at random for each repeat and three experimental repeats are displayed (150 cells total). **D)** Quantifications of cell cycle defects from the single-cell profile plots displayed in C. Bar graphs show mean + SD. **E)** Colony forming assays in RPE1 cells plated at high confluence or with low serum (0.2% FBS) treated with palbociclib (1.25 μM) for 1, 4 or 7 days and then grown at low density without inhibitor for 10 days. Each bar displays mean data + SD from three experiments. Statistical significance determined by Fisher’s exact test (***p < 0.0001). **F)** Cell volume assays in MCF7 and T47D cells treated with 1 μM palbociclib as in A. Graph shows mean data -/+ SD from three repeats. **G)** Western analysis of MCF7 and T47D cells treated as in A. Representative example of 3 repeats.

We hypothesised that the persistent mTOR-dependent growth in cancer lines was driven by oncogenic mutations, which in the case of ER+/HER2- breast cancer cells, are often activating PI3K mutations (PI3K-E545K in MCF7 or PI3K-H1047R in T47D) which signal to mTOR via AKT. In agreement with this hypothesis, inhibiting AKT with the allosteric inhibitor MK-2206 (Hirai et al., 2010) deactivated AKT in all cell types, but only prevented mTOR activity and growth in the breast cancer lines (**Figure 3A-B**). This could also be achieved by reverting the oncogenic PI3K-E545K mutation in MCF7s to wild type (**Figure 3C-D**) (Beaver et al., 2013), which was associated with better cell cycle progression, decreased DNA damage following drug release, and enhanced long-term proliferation (**Figure 3E-H**). Note that most of the replication stress-induced DNA damage occurs after mitosis when chromosomes are incorrectly segregated (Crozier et al., 2022b). Therefore, the fact that oncogene reversion almost doubles the number of cells reaching mitosis (**Figure 3F**) but still reduces overall DNA damage (**Figure 3G**), implies that replication stress is markedly reduced in these cells, most likely because of their restricted growth during G1. Although growth could not be prevented in RPE1 by AKT inhibition, it could be fully suppressed by combined MEK and AKT inhibition, and this was associated with improved cell cycle progression following drug washout (**Figure S2**). This could reflect the dependence of upstream growth factors that stimulate both MEK/PI3K pathways, and/or the oncogenic KRAS mutation present in RPE1 cells (Beaver et al., 2013; Libouban et al., 2017).

**Figure 3:**
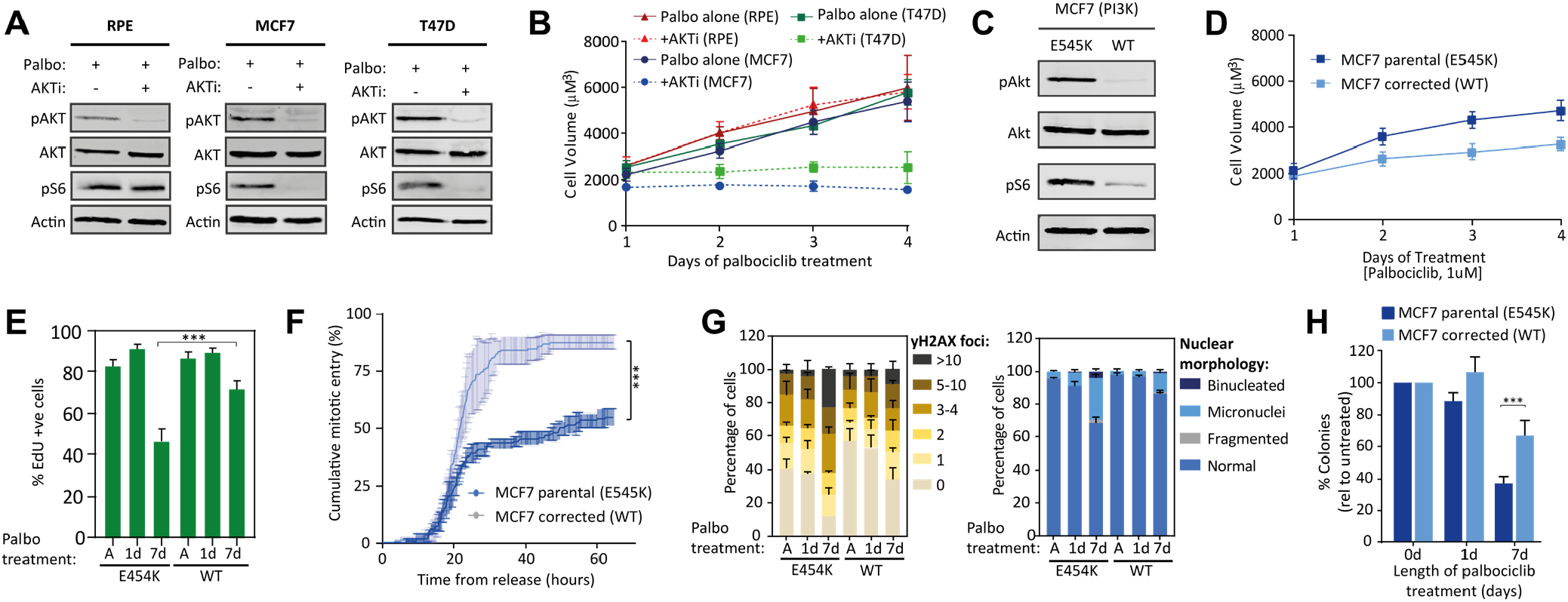
G1 overgrowth and cell cycle exit is driven by oncogenic signals in cancer cells. **A)** Western analysis of indicated cell lines arrested in palbociclib (1.25μM for RPE, 1uM for MCF7/T47D) for 1 day in the presence /absence of the AKT inhibitor (AKTi) MK-2206 (1μM). Representative example of 3 repeats. **B)** Cell volume assays of indicated cell lines arrested in palbociclib (1μM) for 1-4 days in the presence /absence of the AKT inhibitor MK-2206 (1 μM). Graph shows mean data -/+ SD from three repeats. **C)** Western analysis of MCF7 cells, with/without an PI3K-E545K mutation, arrested in palbociclib (1.25μM) for 1 day. Representative example of 3 repeats. **D)** Cell volume assays in MCF7 cells, with/without an PI3K-E545K mutation, arrested in palbociclib (1μM) for 1-4 days. Graph shows mean data -/+ SD from three repeats. **E)** Quantification of the percentage of Edu positive cells following washout from an arrest with 1 μM palbociclib in MCF7 cells, with/without an PI3K-E545K mutation. Cells were treated with DMSO (asynch) or palbociclib for 1 or 7 days, as indicated, and then washed out for 72 hours. EdU was added during the washout period. Data show mean + SD from three experiments, with at least 100 cells quantified per experiment. Statistical significance determined by Fisher’s exact test (*** p < 0.0001). **F)** Cumulative mitotic entry of MCF7 cells, with/without an PI3K-E545K mutation following washout from 7 days treatment with palbociclib (1 μM). A total of 50 cells were quantified per experiment and the graph display mean ± SEM from three experiments. Statistical significance determined by Mann-Whitney test (*** p < 0.0001). **G)** Quantification of γH2AX-positive foci (left panel) and nuclear morphologies (right panel) following palbociclib (1 μM) treatment in MCF7 cells, with/without an PI3K-E545K mutation. Cells were treated with DMSO (asynch) or palbociclib for 1 or 7 days, as indicated, and then analysed after drug washout for 72 h. A total of 100 cells (nuclear morphology) or 50 cells (γH2AX foci) were scored per condition per experiment, and bar graphs represent mean data + SD from three experiments. **H)** Colony forming assays in MCF7 cells, with/without an PI3K-E545K cells treated with palbociclib (1.25 μM) for 1, 4 or 7 days and then grown at low density without inhibitor for 10 days. Each bar displays mean data + SD from 4 experiments. Statistical significance determined by Fisher’s exact test. (*** p < 0.0001).

In summary, oncogenic signals drive excessive growth during a G1 arrest and this leads to DNA damage and long-term cell cycle exit when cells are released from that arrest.

### Oncogenic mutations sensitize non-transformed breast epithelial cells to CDK4/6 inhibition

We next addressed whether non-transformed breast epithelia could be sensitized to CDK4/6 inhibition by introducing oncogenic mutations. We used MCF10A with an endogenous PI3K-E545K or PI3K-H1047R knock-in mutation (Gustin et al., 2009), or MCF10A with a tamoxifen-inducible hRas-G12V mutant (Molina-Arcas et al., 2013). Growth following CDK4/6 inhibition was enhanced in the presence of the oncogene, however this growth plateaued after 2 days in all cells except the hRas cell line, which experienced significant overgrowth during the 4-day treatment (**Figure S3A**). The plateau in net growth was likely related to an inefficient G1 arrest, because all cells, except MCF10A-hRas, were able to continue to proliferate to different extents over the 4-day period of CDK4/6 inhibition (**Figure S3B**). To determine whether this reflected cell cycle delays in all cells, or a penetrant G1 arrest in only a subset of cells, we turned to single cell assays to simultaneously measure cell cycle length (time from mitosis to next mitosis) and cell volume. **Figure 4** demonstrates that all vehicle-treated MCF10A cell lines have a 12h cell cycle length, during which time cell volume increased linearly. The average mitotic volumes and growth rates were not significantly different between any cell type. Following CDK4/6 inhibition, however, cell cycle length was extended in all cells, but this extension was longer in cells expressing oncogenic mutants (**Figure 4A**). The longest cell cycles were observed in hRas expressing cells, which failed to complete a full cell cycle within the 4-day imaging window. A striking effect of these cell cycle delays was that it allowed cells to continue to grow in size. This growth occurred linearly throughout the period of delay and growth rates were not significantly different between the oncogenic mutant lines (**Figure 4B-C**). The net effect was that when cells entered mitosis following a cell cycle delay, they did so with a larger cell volume (**Figure 4D**).

**Figure 4:**
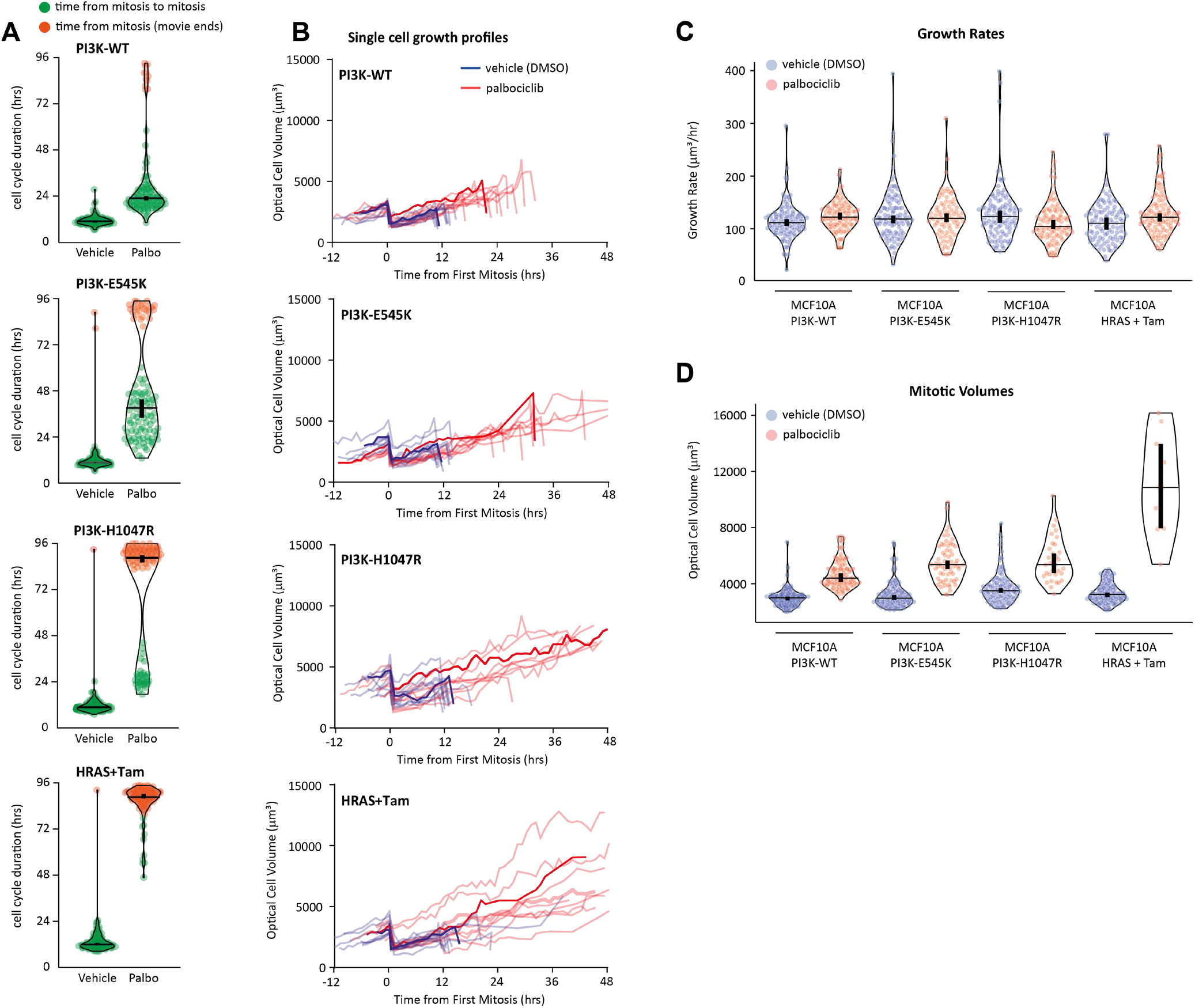
Oncogenes sensitize MCF10A cells to arrest and continue growing following CDK4/6 inhibition. **A-D)** MCF10A cells, or MCF10A expressing different oncogenes, treated with DMSO (vehicle) or palbociclib (1 μM) for up to 4 days and analysed by holographic microscopy to quantify (A) cell cycle duration, (B) single cell optical volume over time, (C) cell growth rates, and (D) optical cell volume during mitosis. In A, C and D, violin plots display data from 100 cells from 2 experiments (except mitotic volumes when some groups has less than 50 mitotic cells per experiment). The thick vertical lines represent a 95% CI around the median (horizontal lines), which can be used for statistical comparison of multiple time points/treatments by eye (see Materials and methods). In B, a typical single cell volume trace is show in blue (vehicle-treated) or red (palbociclib-treated). Random cell volume traces from 10 cells from 2 experiments are show in in light blue/red.

In summary, CDK4/6 inhibition delays cell cycle progression in MCF10A cells and this allows cell to reach a larger size. The oncogene-expressing cells experience longer delays and therefore reach larger overall sizes.

We hypothesised that the overgrown MCF10A cells that experience G1 delays would exhibit DNA damage and p53/p21-dependent cell cycle withdrawal. In agreement, p21 intensity was elevated in cells treated with CDK4/6 inhibitor for 7 days, this was higher in oncogene expressing cells, and this was associated with an inhibition of proliferation after 1-week of treatment (**Figure 5A-B**). In p53-KO cells, p21 induction was prevented and proliferation was rescued over a 3-week treatment period (**Figure 5C-D**). However, cells that continued to proliferate in this condition experienced high levels of DNA damage after only 1 week of treatment (**Figure 5E-H**). Strikingly, MCF10A cells that exhibited slower proliferation in CDK4/6 inhibitor for up to 1 week, could fully recover long-term proliferation when the drug was removed (**Figure 5I**). This indicates that the cell cycle delays and modest increase in size observed in wild type cells (**Figure 4**) does not cause permanent cell cycle exit in this non-transformed epithelial line. This was in sharp contrast to all oncogene expressing MCF10A cells which dramatically lose long-term proliferative potential after as little as 3 days of CDK4/6 inhibitor treatment (**Figure 5I**). Long-term proliferation was partially rescued by p53 knockout (**Figure 5J**), implying that oncogene-dependent cell overgrowth causes p53-dependent cell cycle withdrawal.

**Figure 5:**
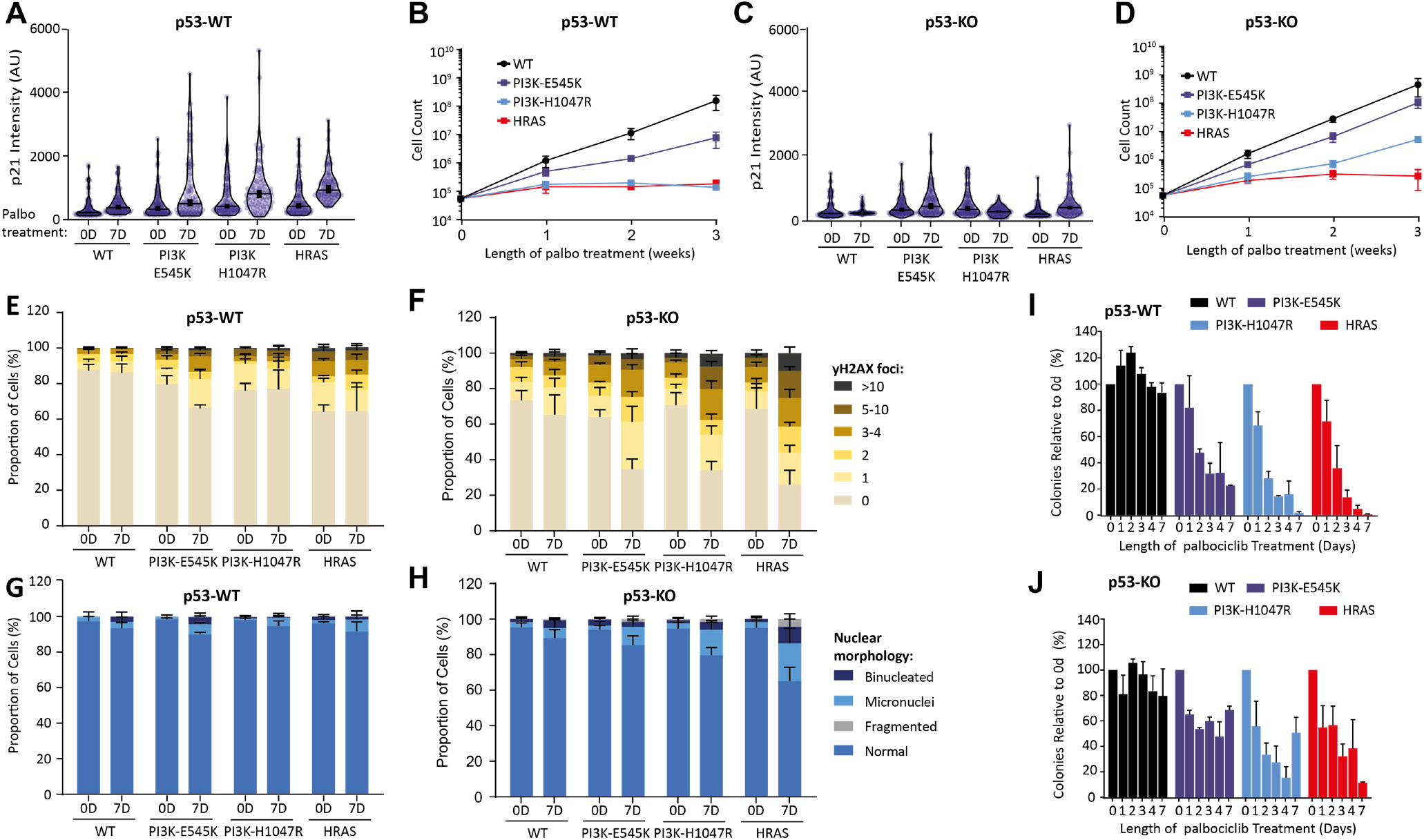
Oncogenes sensitize MCF10A cells to DNA damage and p53/p21-dependent cell cycle exit following CDK4/6 inhibition. **A-D)** p21 intensities (A, C) and cell counts (B, D) in p53-WT and p53-KO MCF10A cells, either WT or expressing different oncogenes, treated continuously in palbociclib (1 μM) for 1-3 weeks, as indicated. A total of 50 cells were scored per condition per experiment from 2-3 experiments. The thick vertical lines in the violin plots represent a 95% CI around the median (horizontal lines), which can be used for statistical comparison of multiple time points/treatments by eye (see Materials and methods). **E-H)** Quantification of γH2AX-positive foci (E-F) and nuclear morphologies (G-H) following palbociclib (1 μM) treatment in MCF10A cells, either WT or expressing different oncogenes. Cells were treated with DMSO (0D) or palbociclib for 7 days (7D), as indicated, and then analysed. A total of 50 cells were scored per condition per experiment, and bar graphs represent mean data + SD from three experiments. **I-J)** Colonies forming assays in p53-WT (I) or P53-KO (J) MCF10A cells, either WT or expressing different oncogenes, treated with palbociclib (1 μM) from 0-7 days, as indicated, and then grown at low density without inhibitor for 10 days. Each bar displays mean data + SD from 3 experiments.

## DISCUSSION

Cell growth must be tightly coupled to cell cycle progression to preserve cell and organismal viability. We demonstrate here that CDK4/6 inhibitors uncouple these two key processes to induce cancer cells to exit the cell cycle. The reason is that oncogenic signals stimulate both cell growth and cell cycle entry, but when the cell cycle is halted in G1 by CDK4/6 inhibition, these oncogenic signals induce continued cell growth that soon becomes toxic. A recent preprint reported similar findings with CDK7 inhibition, implying that different cell cycle drugs may drive senescence via similar mechanisms (Wilson et al., 2021). The concept that hyper-mitogenic signals can drive overgrowth and senescence in arrested cells was proposed nearly two decades ago (Blagosklonny, 2003), but has received little attention since (Blagosklonny, 2022). This is especially surprising since it could explain how general cell cycle inhibitors can produce tumour-specific effects, an age-old problem in cancer research, that has in contrast, received considerable attention over the years. If oncogenes make tumour cells more vulnerable to a G1 cell cycle arrest, then efficiently arresting all cells in G1, for defined periods of time, may lead to cancer-specific overgrowth, DNA damage, and senescence. It will be important compare the rates of growth in different G1-arrested cell types in vivo, because if these differ, then modified dosing schedules may help to optimise cancer-specific overgrowth.

There are many reasons to explain why cell overgrowth is so damaging in G1-arrested cells. Firstly, the overgrowth itself causes gross remodelling of the proteome. Some compartments scale with size, whereas others subscale or superscale (Crozier et al., 2022a; Lanz et al., 2022; Neurohr et al., 2019). Secondly, this atypical scaling induces stress responses that impact on subsequent cell cycle progression. In yeast, G1-arrested cells overgrow and this induces an environmental stress response that is associated with cytoplasmic dilution (Neurohr et al., 2019). In human cells, cytoplasmic dilution may also occur following CDK4/6 inhibition leading to decreased macromolecular crowding (Neurohr et al., 2019). Consistent with this model, Crozier et al demonstrate that overgrowth triggers an osmotic stress response that is associated with increased intracellular osmolyte concentrations (Crozier et al., 2022a). This osmotic stress response causes p21 induction via a p38-mediated pathway, resulting in delayed and attenuated exit from G1 when CDK4/6 inhibitors are removed. Thirdly, some cells can escape this G1 arrest, but these cells experience significant replication stress during the proceeding S-phase, which is associated with dilution of replisome components that subscale with size (Crozier et al., 2022a; Crozier et al., 2022b). Very recent data suggest that the DNA damage response is also impaired in enlarged cells, and the DNA itself may be prone to damage (Manohar et al., 2022). As a result of these replication-associated problems, p21 protein is induced again as cells enter G2, causing further cell cycle exit (Crozier et al., 2022a). Finally, cells that fail to exit the cell cycle from G2, in particular p53-deficient cells that cannot induce p21, enter mitosis and experience even further damage due to catastrophic chromosome segregation errors, promoting further cell cycle withdrawals (Crozier et al., 2022a; Crozier et al., 2022b; Manohar et al., 2022; Wang et al., 2022). Future research will be important to determine how much these various routes to genotoxic stress contribute to cell cycle exit in cancer cells with different oncogenic mutations.

It is now crucial to validate the findings we demonstrate here in animal models and patient samples, because if oncogene-dependent cell overgrowth is important to drive DNA damage and cell cycle exit in vivo then this would have important clinical implications. Most importantly, it would rationalise effective combination therapies that converge to inhibit Cyclin D-CDK4/6 activity *without* affecting global translation rates. Interestingly, this is the predicted effect of anti-estrogen therapy in combination with CDK4/6 inhibitors, which is the current standard-of-care treatment in HR+/HER2- breast cancer (Fassl et al., 2022). That is because estrogens stimulate estrogen receptors to translocate to the nucleus and enhance the transcription of Cyclin D (Prall et al., 1997; Sabbah et al., 1999). Analogous synthetic lethal combinations have been proposed in clear cell renal cell carcinoma, where HIF2A stimulates Cyclin D transcription (Nicholson et al., 2019). Furthermore, other combinations have been identified that act via similar principles, such as CK1ε inhibition which represses SP1-mediated CDK6 transcription following CDK4/6 inhibition in breast cancer (Dang et al., 2021). The ability of cancer cells to transcriptionally upregulate Cyclin D-CDK4/6 activity via a variety of different routes, suggests that similar effective combinations await discovery in other tumour types.

Multiple clinical trials are currently testing inhibitors of growth factor signalling pathways, such as PI3K, MEK, AKT and mTOR, together with CDK4/6 inhibitors (Fassl et al., 2022). The principle of using these combinations to achieve a sensitized G1 arrest has been extensively demonstrated in preclinical models, therefore it is logical to assume that a similar sensitized G1 arrest will also be beneficial for treating patient tumours (Alvarez-Fernandez and Malumbres, 2020; Watt and Goel, 2022). Genomic alterations that are predicted to enhance growth factor signalling pathways are also frequently associated with resistance to CDK4/6 inhibition in patients (Costa et al., 2020; Formisano et al., 2019; Wander et al., 2020), which further underscores the benefit of inhibiting these pathways in combination to effectively block G1 progression. Nevertheless, the question of whether the resulting G1 arrest is cytostatic or cytotoxic is still likely to be crucial for the overall response and this will depend on the mechanism(s) used to clear G1-arrested tumour cells in vivo. If this is via overgrowth-mediated cell cycle withdrawal, as we show here, then the response will likely depend on the extent and type of overgrowth, which will be defined by the oncogene(s) driving that growth and the particular drug combination used to arrest cells in G1. It is important to point out however, that combined inhibition of CDK4/6 and growth factor pathways can lead to either senescence or apoptosis in preclinical models (Wagner and Gil, 2020), therefore which endpoint is more clinically-relevant is debated. Some studies demonstrate that G1-arrested tumour cells can die via apoptosis following combined inhibition of CDK4/6 and growth factor signalling (Herrera-Abreu et al., 2016; Jansen et al., 2017; Zhao et al., 2021), whereas other studies show that cytostasis or senescence is more prevalent (Formisano et al., 2019; Goel et al., 2016; Michaloglou et al., 2018; Vora et al., 2014). It can be challenging to monitor senescence in clinical samples, but this is an important future goal, as discussed in this review (Witkiewicz et al., 2022). At this stage, we would simply urge the monitoring of cell size and DNA damage in proliferating tumour cells, before and during treatment, to assess whether these properties change and, if so, whether that is predictive of the overall response. If increased cell size correlates with a positive response, then this may help to define predictive biomarkers since mTOR activity and TSC1/2 status varies in HR+/HER2- breast cancer lines, and this has been shown to causes differential sensitivity to CDK4/6 inhibition (Maskey et al., 2021).

Another major finding of this work is that oncogenes can drive cell cycle delays, cell overgrowth and DNA damage in MCF10A cells treated continually with CDK4/6 inhibitor. This leads to p53-dependent p21 activation and cell cycle exit over time. Similar cell cycle delays and DNA damage were observed in a range of p53-deficient cancer lines treated continuously with CDK4/6 inhibitor (Crozier et al., 2022b). Therefore, continual CDK4/6 inhibition, if it does not lead to a penetrant G1 arrest immediately, can still lead to cell size deregulation, DNA damage and p53-dependent cell cycle withdrawal over time. This may explain why p53 is associated with resistance in patients (Patnaik et al., 2016; Wander et al., 2020), and it could open new possibilities for using CDK4/6 inhibitors to tackle hard-to-treat cancers that lack p53. CDK4/6 inhibition may not drive cell cycle exit directly in these cells, but if the tumour cells become enlarged and damaged following treatment then this could produce vulnerabilities to secondary agents, such as chemotherapeutics that either enhance DNA damage or inhibit DNA damage repair. This may help to explain the surprising finding that the CDK4/6 inhibitor trilaciclib improves overall survival in triple-negative breast cancer when given prior to chemotherapy with the genotoxic combination of gemcitabine and carboplatin (Tan et al., 2022).

In summary, we demonstrate here that CDK4/6 inhibitors allow oncogenic signals to drive toxic cell overgrowth and cell cycle exit, which holds great promise for the long-term goal of achieving tumour-selectivity with general cell cycle inhibitors. It is now crucial to validate these findings in animal models and patient samples, because if this is also observed in vivo, then this ability of CDK4/6 inhibitors to switch pro-proliferative oncogenic signals into toxic anti-proliferative responses would represent a new paradigm for anti-cancer treatment. Tumours may be addicted to oncogenes for survival, but if that addiction could be turned into a liability by inhibiting the cell cycle, then oncogenes could yet prove to be cancer’s Achilles heel.

## ACKNOWLEDGMENTS

We thank the Dundee Imaging Facility and Genetic Core Services for help with this work, and Tony Ly and Alexis Barr for critical reading of this manuscript. This work was supported by Tenovus Scotland (studentship for RF), a Cancer Research UK Programme Foundation Award to ATS (C47320/A21229. which also funds LC), and the Ninewells Cancer Campaign (studentship for AP).

## AUTHOR CONTRIBUTIONS

A.T. S., R.F. and L.C. conceived the study, designed the experiments, and interpreted the data. R.F. and L.C. performed majority of experiments with important contributions from A.P. B.H.P. provided the MCF7 and MCF10A PI3K knock-in cell lines. A.T. S. supervised the study and wrote the manuscript with help from R.F, L.C. and A.P.

## MATERIALS AND METHODS

### Cell culture and reagent

hTERT-RPE1 (RPE1) cells were purchased from ATCC and the RPE1-FUCCI were published previously (Krenning et al., 2014). The human ER+/HER2- breast cancer lines, MCF7 and T47D, were purchased from ATCC. The MCF7 parental (PI3KA-E545K) and corrected (PI3KA-WT) cell lines are described in (Beaver et al., 2013). The MCF10A PI3K-E545K and H1047R knock-in lines are described in (Gustin et al., 2009) and the inducible hRAS lines is described in (Molina-Arcas et al., 2013). P53 knockout cell lines were generated by CRISPR/Cas9 using a gRNA targeting exon 4 of p53 (ACCAGCAGCTCCTACACCGG) followed by selection in 5μM Nutlin-3A, as described previously (Crozier et al., 2022). All cells were authenticated by STR profiling (Eurofins) and screened for mycoplasma every 1–2 months. RPE, MCF7 and T47D cells were cultured at 37°C with 5% CO_2_ in DMEM (Thermo Fisher Scientific, Gibco 41966029) supplemented with 9% FBS (Thermo Fisher Scientific, Gibco 10270106) and 50 μg/ml penicillin/streptomycin (Sigma, P4458). MCF10A cells were cultured in F12/DMEM (Thermo Fischer Scientific, Gibco, 11320033) and supplemented with 5% horse serum (Thermo Fischer Scientific, Gibco 16050122), 20ng/mg EGF (Sigma, E9644), 0.5ug/ml hydrocortisone (Sigma, H088), 100ng/ml cholera toxin (Sigma, C8052), 10ug/ml insulin (Sigma, I9278) and 50ug/ml penicillin/streptomycin (Sigma, P4458). The following drugs were used in this study: Palbociclib (PD-0332991, hydrochloride salt, MedChemExpress, HY-50767A), EdU (Sigma-Aldrich, BCK-EDU488). Gedatolisib (PF-05212384, Sigma, PZ0281), Mirdametinib (PD-0325901, Selleckchem, S1036), MK-2206 (MedChemExpress, HY-108232), nutlin-3a (Sigma, SML0580), S-Trityl-L-cysteine (STLC; Sigma Aldrich, #164739), DAPI (4′,6-Diamidino-2-Phenylindole; Thermo Fisher Scientific, D1306),

### Cell density

To prevent cell–cell contact from inhibiting exit from G1, it was crucial to plate cells at low density for all experiments, except those plated at high confluence shown in fig.2 (in this case cells were plated at 100% confluence). This is especially true for RPE1 cells that arrest the cell cycle upon contact inhibition. Therefore, cells were plated at a maximum density of 8,000 cells per cm2 immediately prior to the arrest with CDK4/6 inhibitors.

### Cell volume measurements

For volume measurement, cells were plated in 6-well plates with palbociclib -/+ other treatments for 1-4 days. Washed and trypsinised cells were analysed on an NC-3000 Nucleocounter to quantify diameter using DAPI and Acridine orange. The histograms containing information of cell diameter was imported to Flowing Software version 5.2.1 and the appropriate gates were added to include the main peak of the histogram. Cell volume was then calculated as 4/3 πr^3^.

### Determining protein and RNA concentration

For protein and RNA concentration measurements, cells were plated in 10cm dishes with 1μM palbociclib for 1-7 days, after which sells were harvested and protein concertation calculated with a detergent compatible (DC) assay (BioRad), or RNA concentration determined using a Trizol-based method (Thermo Fisher Scientific, Invitrogen 15596018).

### Immunofluorescence

Cells were plated at low density on High Precision 1.5H 12-mm coverslips (Marienfeld) and fixed for 10 min with 4% paraformaldehyde dissolved in PBS. Once fixed, coverslips were washed three times in PBS and then blocked in 3% BSA dissolved in PBS with 0.5% Triton X-100 for 30 min. Coverslips were then incubated with primary antibodies at 4°C overnight, prior to washing with PBS and incubation with secondary antibodies and DAPI (1 μg/ml) for 2–4 h at room temperature. After further washing, coverslips were mounted onto slides with ProLong Gold Antifade (Thermo Fisher Scientific, P10144). Coverslips were imaged on either a Zeiss Axio Observer using a Plan-apochromat 20×/0.8 M27 Air objective or a Deltavision with a 100×/1.40 NA U Plan S Apochromat objective. The primary antibodies used were as follows: mouse anti-phospho-Histone H2A.X (Ser139; clone JBW301; Sigma, 05-636; 1/1,000), mouse tubulin (clone B-5-1-2, Sigma, T5168-.2ML; 1/5000), and p21 Waf1/Cip1 (clone 12D1, Cell Signaling Technology, #2947, 1:1000). The secondary antibodies used were highly cross-absorbed goat anti-rabbit or anti-mouse coupled to Alexa Fluor 488 and Alexa Fluor 568 which were all used at 1/1,000 dilution. All antibodies were made up in 3% BSA in PBS. For EdU staining, a base click EdU staining kit was used (Sigma, BCK-EDU488), as per manufacturer’s instructions.

### Time-lapse imaging

For FUCCI time-lapse imaging, cells were plated at low density (approximately 15,000 cells per well) and imaged in 24-well plates in DMEM inside a heated 37°C chamber with 5% CO2. Images were taken every 10 min with a 10×/0.5 NA air objective using a Zeiss Axio Observer 7 with a CMOS ORCA flash 4.0 camera at 4 × 4 binning. For bright-field imaging, cells were imaged in a 24-well plate in DMEM in a heated chamber (37°C and 5% CO2) with a 10×/0.5 NA air objective using a Hamamatsu ORCA-ER camera at 2 × 2 binning on a Zeiss Axiovert 200 M, controlled by Micro-manager software (open source; https://micro-manager.org/) or with a 20×/0.4 NA air objective using a Zeiss Axio Observer 7 (details above). Holographic imaging movies were capture using a Holomonitor M4 microscope to quantify single-cell volumes, which were calculated using the associated software.

### Image analysis, quantification, and statistics

All holomonitor analysis was carried out with a Holomonitor M4. A total of 15,000 cells were plated into either a Sarstedt lumox multiwell, or an Ibidi μ-plate glass-bottomed 24 well plate. After 24hrs cells were treated and imaging began immediately. Images were scheduled every 20 minutes for a total of 96 hours. For single cell traces 5 cells were randomly selected in the first frame then followed by eye with volume measurements being taken every 2 hours for the duration of the first full cell cycle (mitosis to mitosis) or until the 48hr mark. For population analysis 50 cells were randomly selected and the volume and time of their first and second mitosis following treatment was recorded. Growth rates were assumed to be linear and were calculated as (V2-(V1-2))/CL, where V1 is equal to the volume of the first mitosis, V2 is equal to the volume of the second mitosis, and CL is equal to the time between the first and second mitosis. For cells that did not enter mitosis a second time during the imaging period, V2 was taken as interphase volume 48hrs after first mitosis, so growth rate could still be calculated.

The single-cell FUCCI profiles were generated manually by analysing RPE1-FUCCI movies. A total of 50 red cells were randomly selected and marked at the beginning of the movie. The time points in which the FUCCI cells change colour was recorded to determine the time spent in each phase of the first cell cycle following release from CDK4/6 inhibition. All images were placed on the same scale prior to analysis to ensure that the red/yellow/green cut-offs were reproducibly calculated between experiments, which we performed using identical illumination conditions. Mitotic entry was timed based on the visualization of typical mitotic cell rounding and loss of nuclear mAG-geminin signal. For mitotic entry quantifications in brightfield movies, 50 cells were selected at random at the beginning of the time lapse and the time point that cells entered mitosis was determined. Mitotic entry was timed based on when the nuclear envelop breaks down and the cell rounds up. γH2AX foci were counted by eye in the first 50 cells (per condition) selected using the DAPI channel. For scoring of nuclear abnormalities, the first 100 cells within the image were counted and scored based on their nuclear morphology. To quantify EdU incorporation 100 cells were randomly selected in the DAPI channel and the number of EdU-positive cells was then counted.

Statistical significance was determined by Fisher’s exact test or Mann-Whitney tests, as indicated in legends. The graphs in Figs 4 and Fig.5a, 5c are plotted as violin plots using PlotsOfData (Postma and Goedhart, 2019); https://huygens.science.uva.nl/PlotsOfData. This allows the spread of data to be accurately visualized along with the 95% confidence intervals (thick vertical bars) calculated around the median (thin horizontal lines). Statistical comparison can then be made by eye between any treatment and time points, because when the vertical bar of one condition does not overlap with one in another condition, the difference between the medians is statistically significant (P < 0.05).

### Western blotting

Total protein lysates for immunoblot were prepared by scraping cells into 4X protein loading buffer (250mM Tris, 10% SDS, 40% Glycerol, 0.1% Bromophenol Blue) and then sonicating with a Cole-Parmer ultrasonic processor (20% Amp, 15 sec pulse). Samples were briefly boiled and centrifuged followed by a DC assay to determine protein concentration, after which 2-mercaptoethanol was added at a final concentration of 10%. Equal concentrations of protein were loaded and then separated on SDS—PAGE gels and transferred to 0.45μm nitrocellulose membranes (Amersham Protran Premium). After transfer, blots were blocked in 5% milk in TBS with 0.1% Tween 20 (TBS-T) and incubated overnight at 4°C in primary antibody in TBS-T. Membranes were then washed three times in TBS-T, incubated in IRDye secondary antibody for 2h, and washed 3 further times prior to visualisation on a LI-COR Odyssey CLx system. The primary antibodies used were rabbit pS6 (Phospho-S6 Ribosomal Protein; Ser235/236, Cell Signalling, 4858, 1/1000), rabbit actin (Sigma, A2066, 1/5000) rabbit pAKT (Ser473, Cell Signalling, 4060B, 1/1000), rabbit AKT (Cell Signalling, 9272, 1/1000), rabbit ERK1/2 (Upstate, 06-182, 1/1000), mouse pERK1/2 (Thr202/Try204, Cell Signalling, 9106S), 1/1000). Secondary antibodies used were IRDye 800CW Goat anti-Mouse IgG (LI-COR) or IRDye 800CW Goat anti-Rabbit IgG (LI-COR). Both LI-COR secondary antibodies were used at a 1/15,000 dilution.

### Colony-forming assays

For the colony-forming assays, cells were treated with palbociclib at 60,000 cells per 10 cm dish for different length of time (1–7 days) prior to drug washout (6 × 1 h washes). Following washing and trypsinization, RPE1 and MCF10As were plated in triplicate at 250 cells into 10 cm dishes and left to grow for 10 days, whereas MCF7 and T47Ds were plated at 500 cells in triplicate in 6-well plates and allowed to grow for 14–21 days. At the end of the assay, cells were washed twice in PBS and then fixed at 100% ethanol for 5 min. Developing solution (1:1 ratio of 2% Borax:2% Toluene-D in water) was added to the fixed cells for 5 min and the plates were then rinsed thoroughly with water and left to dry overnight. The plates were then scanned and the number of colonies were quantified using ImageJ. This was performed by cropping to an individual plate and converting to a binary image. The fill holes, watershed and analyse particles functions were then used to count colonies.

### Weekly fold increase in cell count

A total of 60,000 cells from each MCF10A line were plated into 10 cm dishes and treated with palbociclib (1μM) or DMSO (control). After 7 days of treatment, cells were trypsinized, total cell counts were determined and the 7-day fold increase in cell count was calculated. From the cell suspension, 60,000 cells were returned to palbociclib treatment, and this process was repeated two more times for a total of 3 weeks. At each time point, excess cells were transferred to coverslips and taken for immunofluorescence with γH2AX antibodies

