## Supplementary Figures for "Oncogenic signals prime cancer cells for toxic cell growth during a G1 cell cycle arrest"

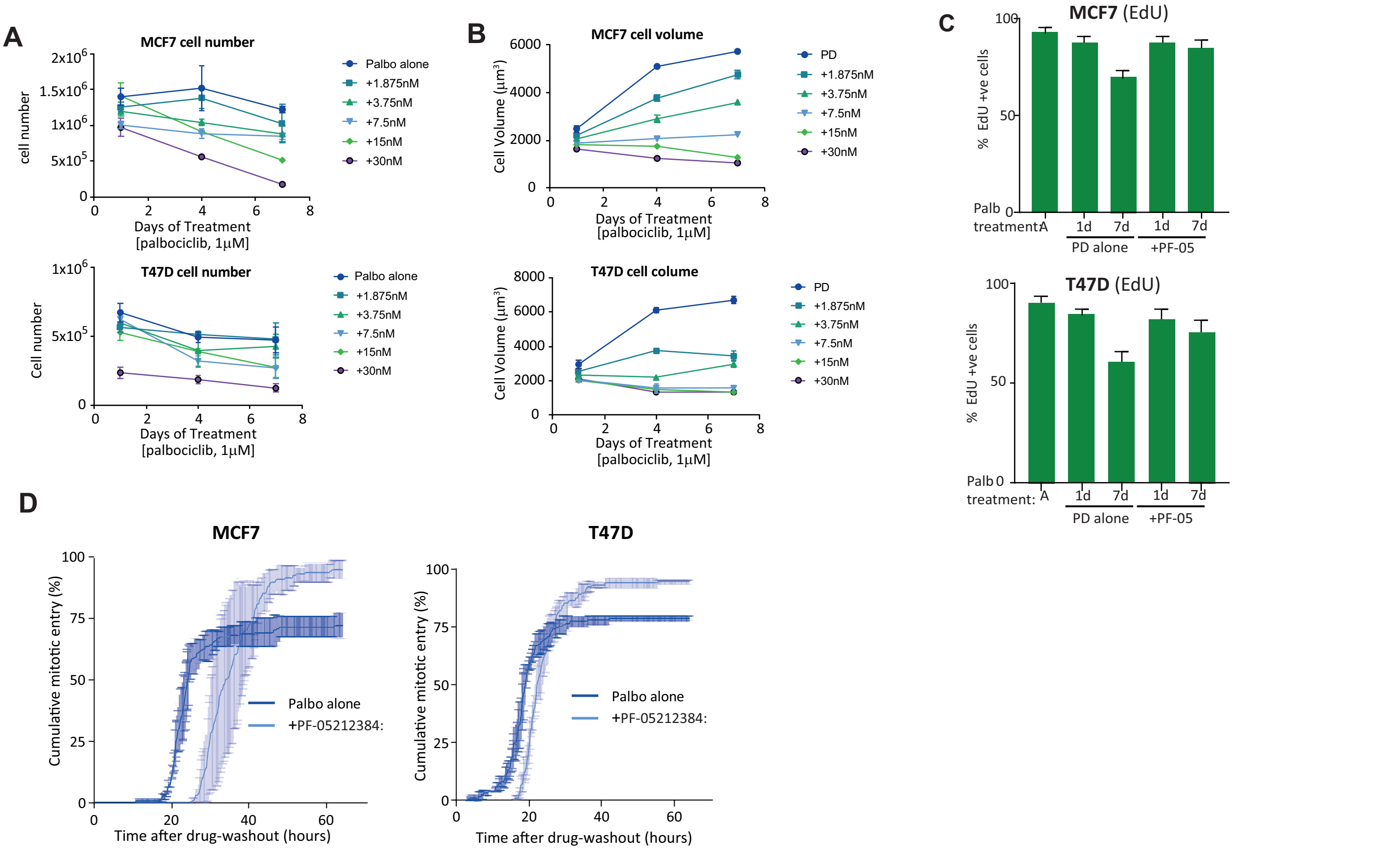


**Figure S1: PF-05212384 reduces growth during G1, and enhances cell cycle re-entry and long-term proliferation in breast cancer cells. A-B)** Quantification of cell number (A) and cell volume (B) of MCF7 or T47D cells arrested in palbociclib (1 μM) for 1-4 days in the absence or presence PF-05212384 at indicated concentrations. Graph shows mean data from three repeats. Graph shows mean data -/+ SD from three repeats. **C)** Quantification of the percentage of Edu positive cells following washout from an arrest with 1 μM palbociclib in MCF7/T47D cells -/+ PF-05212384 (7.5 nM). Cells were treated with DMSO (asynch) or palbociclib (1 μM) for 1 or 7 days then washed out for 72 hours. EdU was added during the washout period. Data show mean + SD from three experiments. **D)** Cumulative mitotic entry in MCF7 (top panel) or T47D (bottom panel) following washout from 7 days treated of palbociclib (1 μM) +/- PF-05212384 (7.5 nM). A total of 50 cells were quantified per experiments and graphs display mean ± SEM from three experiments.

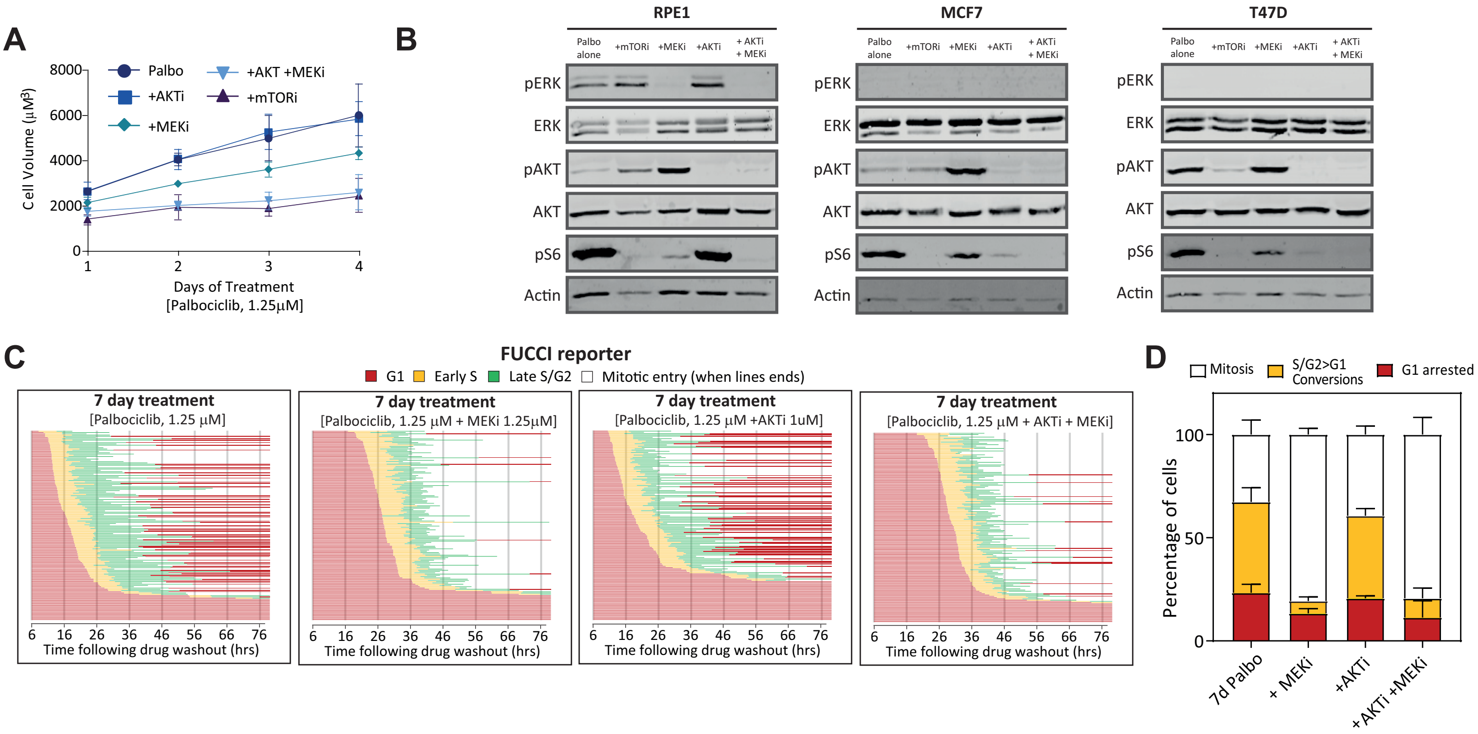


**Figure S2: Combined MEK and AKT inhibition on G1 growth and cell cycle exit in RPE1 cells. A)** Volume assays of RPE1 cells arrested in palbociclib (1.25μM) for 1-4 days either alone, with 1.25μM PD-0325901 (MEKi), with 1 μM MK-2206 (AKTi), or with 1 μM MK-2206 and 1.25μM PD-0325901 combined (AKTi + MEKi). Graph shows mean data -/+ SD from three repeats. **B)** Western analysis of RPEs, MCF7 or T47D cells arrested in palbociclib (1.25 μM RPE1, 1 μM MCF7/T47F) for 1 days either alone, or with 30nM PF-05212384 (mTORi), 1.25 μM PD-0325901 (MEKi), 1 μM MK-2206 (AKTi), or 1 μM MK-2206 and 1.25 μM PD-0325901 combined (AKTi + MEKi). Representative example of 1 repeat. **C)** Cell cycle profile of individual RPE1-FUCCI cells (each bar represents one cell) after washout from 1 or 7 days of treatment with palbociclib (1.25 μM) +/- 1.25 μM PD-0325901 (MEKi), 1 μM MK-2206 (AKTi), or 1 μM MK-2206 and 1.25 μM PD-0325901 combined (AKTi + MEKi). STLC (10 μM) was added to prevent progression past the first mitosis. Fifty cells were analysed at random for each repeat and three experimental repeats are displayed (150 cells total). **D)** Quantifications of cell cycle defects from the displayed single-cell profile plots. Bar graphs show mean + SD.

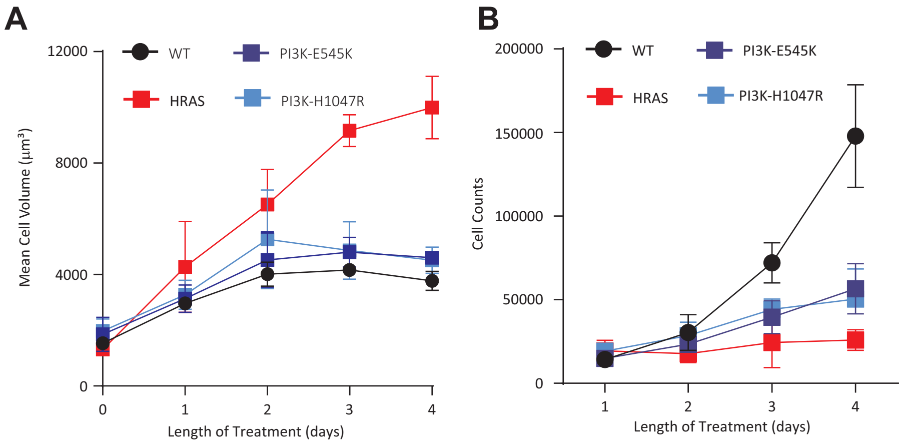


**Figure S3: Incomplete arrest in MCF10A lines after CDK4/6 inhibition. A-B)** Mean cell volume (A) and cell counts (B) in MCF10A cells, either WT or expressing different oncogenes, treated continuously in palbociclib (1 μM) for 1-4 days, as indicated. Graphs shows mean data -/+ SD from three repeats.
